# Selectively increased autofluorescence at fingernails and certain regions of skin: A potential novel diagnostic biomarker for Parkinson’s disease

**DOI:** 10.1101/322222

**Authors:** Danhong Wu, Yue Tao, Mingchao Zhang, Yujia Li, Liwei Shen, Yuzhen Li, Weihai Ying

## Abstract

Diagnosis of Parkinson’s disease (PD) mainly relies on the judgment of experienced neurologists on the clinical symptoms of patients. Quantitative and specific biomarker tests for PD is greatly needed. In this study we tested our hypothesis that increased autofluorescence (AF) of skin and fingernails may become a novel diagnostic biomarker for PD. Our study has indicated that PD patients have a distinct pattern of AF changes, compared with that of acute ischemic stroke (AIS) patients: First, the AF intensity of PD patients in the fingernails and a part of the examined regions of skin is significantly higher than that of the healthy and Low-Risk group, while the AF intensity of AIS patients is significantly higher than that of the healthy and Low-Risk group in most regions examined; second, there is AF asymmetry at the index fingernails and two regions of the skin of PD patients, while there is AF asymmetry at all examined regions of AIS patients; and third, both the AF intensity and AF asymmetry at Centremetacarpus of PD patients is significantly lower than those of AIS patients. The increased AF may result from the altered keratins’ AF induced by the oxidative stress in the plasma of PD patients. Collectively, our study has indicated that PD patients have a distinct pattern of AF changes compared with those of healthy and Low-Risk persons as well as AIS patients, which may become a novel diagnostic biomarker for PD.

## Introduction

Parkinson’s disease (PD) is the second most common neurodegenerative disorder characterized by resting tremor, muscular rigidity and gait disturbances (7). PD pathology is characterized by the progressive loss of dopaminergic neurons in the substantia nigra (SN) pars compacta and their termini in the dorsal striatum (17), with the presence of Lewy bodies - the intracellular inclusions of aggregated α-synuclein - as the hallmark of the disorder (16). The pathological mechanisms of PD include neuroinflammation, oxidative stress, mitochondrial dysfunction, and abnormal processing of misfolded proteins (3,8,9,18). The prevalence of the disorder is approximately 1 - 2% in people over age 60 in industrialized countries. In 2015, PD affected 6.2 million people and resulted in approximately 117,000 deaths globally (5,14). It is projected that the number of individuals with PD in the world will triple by 2030 (15).

There are several major problems and challenges in current PD diagnosis: 1) Diagnostic criteria for PD are nonspecific clinical symptoms of patients, including rest tremor, bradykinesia and cogwheel rigidity, while the gold standard for PD remains to be neuropathologic confirmation. The accuracy and efficacy of PD diagnosis has been limited by the lack of biomarker tests (1). 2) PD diagnosis mainly relies on the clinical judgment of experienced neurologists. Therefore, the shortage of experienced neurologists in many areas may lead to significant misdiagnosis for PD. 3) The diagnostic accuracy for early-stage PD has been particularly unsatisfactory: It was reported that clinical diagnosis of PD in untreated or not clearly responsive subjects had only 26% accuracy (1). Therefore, one of the critical requirements for effective PD treatment is to establish new approaches for accurate and efficient PD diagnosis.

Human autofluorescence (AF) has been used for non-invasive diagnosis of diabetes, which is based on detection of the AF of advanced glycation end-products (AGEs) of the collagen in dermis (11,13). It is of significance to further investigate the potential of AF as biomarkers for diseases, due to the non-invasiveness and simplicity of AF detection. Our recent study has suggested that UV-induced epidermal green AF, which is originated from UV-induced keratin 1 proteolysis in the spinous cells of epidermis, is the first non-invasive biomarker for predicting UV-induced skin damage (10). We have further found that the oxidative stress induced by UVC is causative to the increased epidermal AF of mouse ears by inducing keratin 1 proteolysis (12). Since there is increased oxidative stress in the plasma of PD patients (2,4,6), it is warranted to determine if there are increases in the epidermal AF of PD patients, which may become a novel diagnostic biomarker for the disease.

In this study, we tested our hypothesis that PD patients may have increased AF in fingernails and certain regions of their skin. Our study has provided first evidence that there are selective increases in green AF intensity in the fingernails and certain regions of the skin of PD patients, which has provided evidence supporting our hypothesis.

## Methods

### Human subjects

The study was conducted according to a protocol approved by the Ethics Committee of Shanghai Fifth People’s Hospital, Fudan University. The human subjects in our study were divided into three groups: Group 1: The group of healthy persons and Low-Risk persons - the persons with only a mild level of hypertension; Group 2: The group of PD patients - the people who were clinically diagnosed as PD by the neurologists of the Department of Neurology, Shanghai Fifth People’s Hospital, Fudan University; and Group 3: The group of AIS patients who had ischemic stroke within 7 days from the time of examination. The age of Group 1, Group 2 and Group 3 is 67.44 ± 7.24, 72.9 ± 7.91, and 67.30 ± 10.27 years of old, respectively.

### Autofluorescence determination

A portable AF imaging equipment was used to detect the AF of the fingernails and certain regions of the skin of the human subjects. The excitation wavelength is 485 nm, and the emission wavelength is 500 - 550 nm. For all of the human subjects, the AF intensity in the following six regions on both hands, i.e., twelve regions in total, was determined, including the index fingernails, Ventroforefingers, dorsal index fingers, Centremetacarpus, Ventriantebrachium, and Dorsal Antebrachium.

### Statistical analyses

All data are presented as mean ± SEM. Data were assessed by one-way ANOVA, followed by Student - Newman - Keuls *post hoc* test, except where noted. *P* values less than 0.05 were considered statistically significant.

## Results

We determined the green AF intensity of Dorsal Antebrachium of Healthy and Low-Risk group, the PD group, and the AIS group. On right and left Dorsal Antebrachium, both PD patients and AIS patients have significantly higher green AF intensity than Healthy and Low-Risk persons (Fig. 1A). We further determined the AF asymmetry at the Dorsal Antebrachium of these three groups, showing that both PD patients and AIS patients have significantly higher AF asymmetry than Healthy and Low-Risk persons (Fig. 1B). On right and left Ventroforefinger, both PD patients and AIS patients also have significantly higher green AF intensity than Healthy and Low-Risk persons (Fig. 2A). The AIS patients have significantly higher AF asymmetry than Healthy and Low-Risk persons (Fig. 2B).

**Fig. 1.**
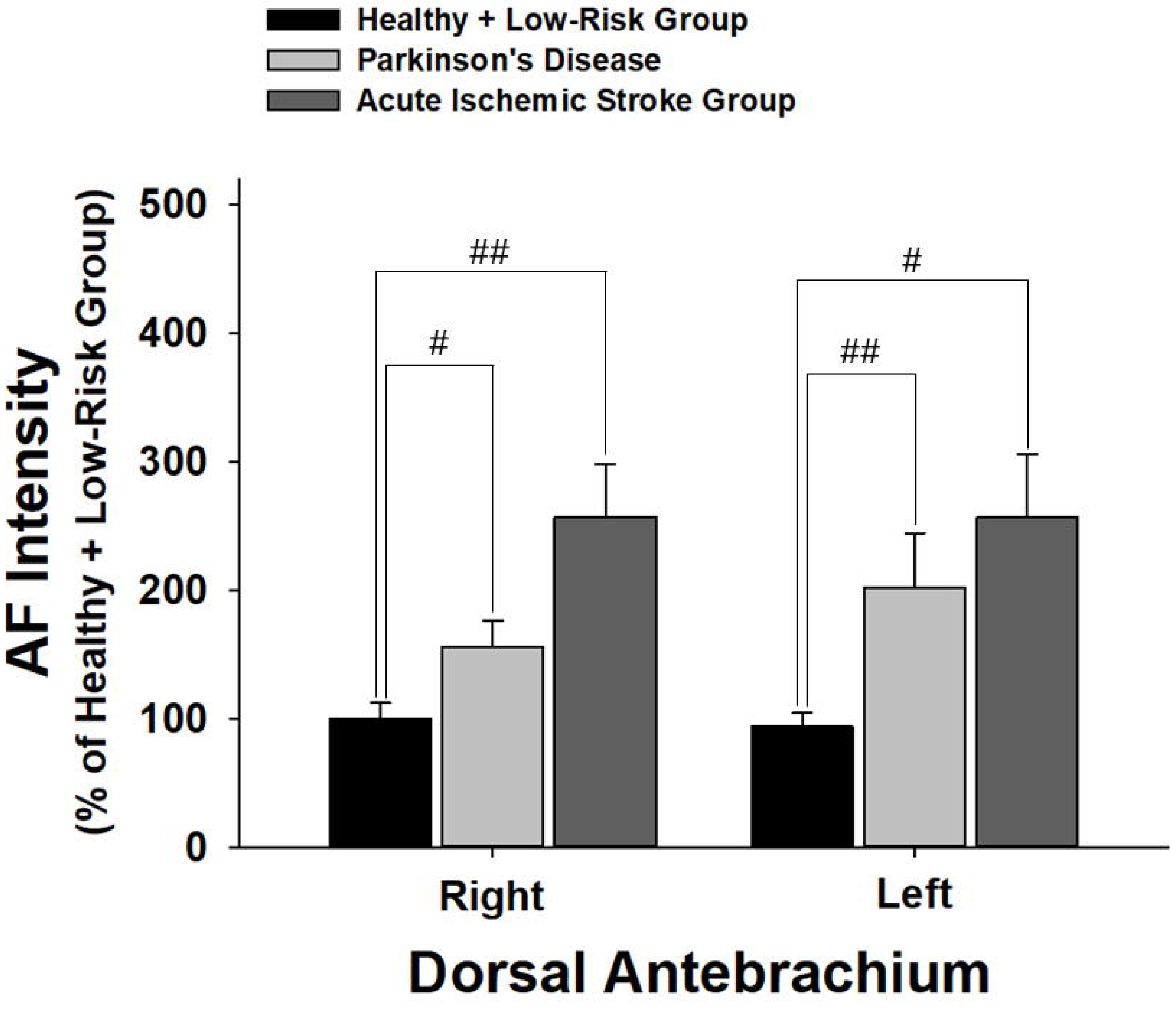
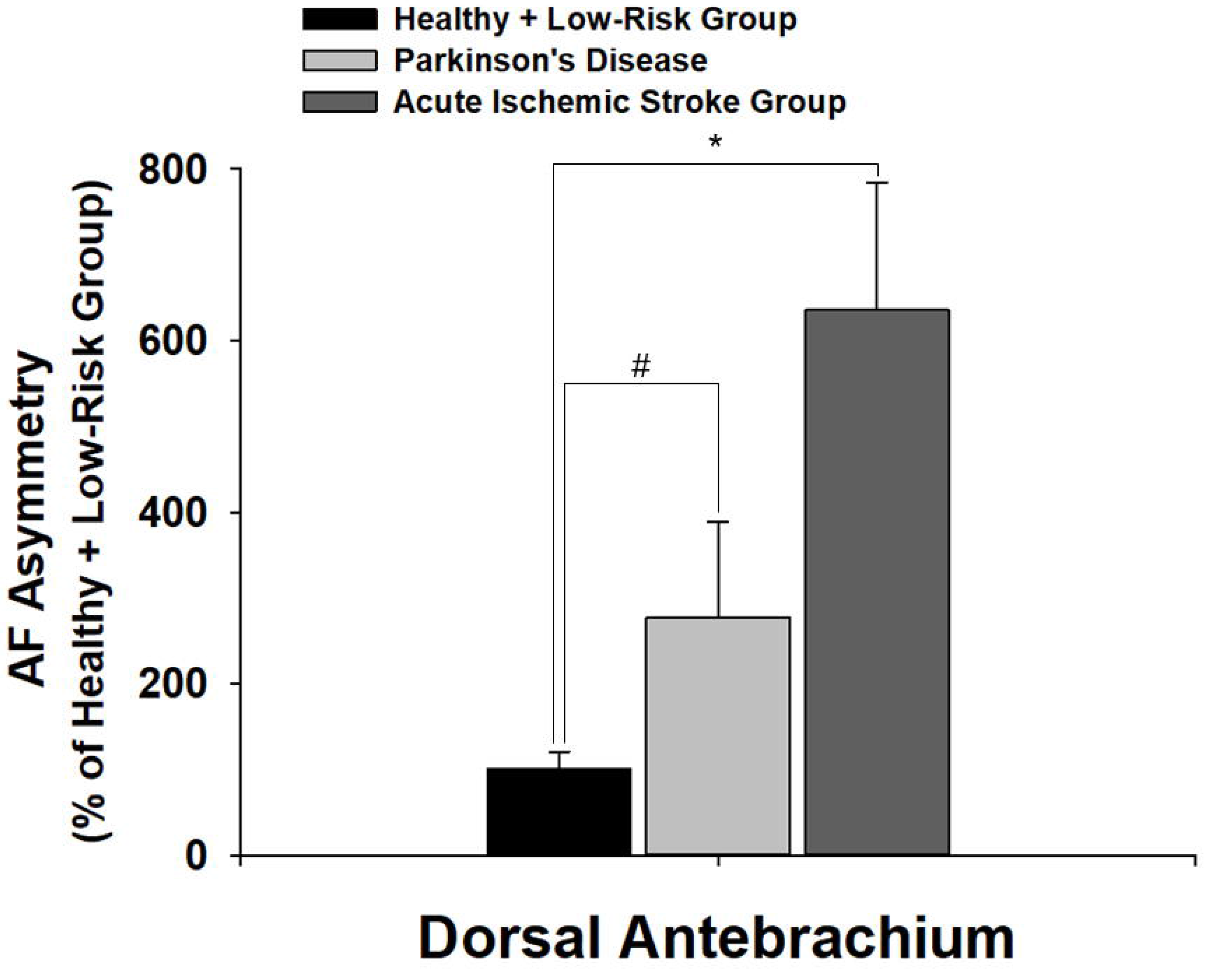
(A) On right and left Dorsal Antebrachium, both PD patients and AIS patients have significantly higher green AF intensity than Healthy and Low-Risk persons. (B) Both PD patients and AIS patients have significantly higher AF asymmetry than Healthy and Low-Risk persons at Dorsal Antebrachium. The number of subjects in the Healthy and Low-Risk group, the PD group, and the AIS group is 25, 13, and 47-49, respectively. *, *p* < 0.05; #, *p* < 0.05 (Student t-test); ##, *p* < 0.01 (Student *t*-test).

**Fig. 2.**
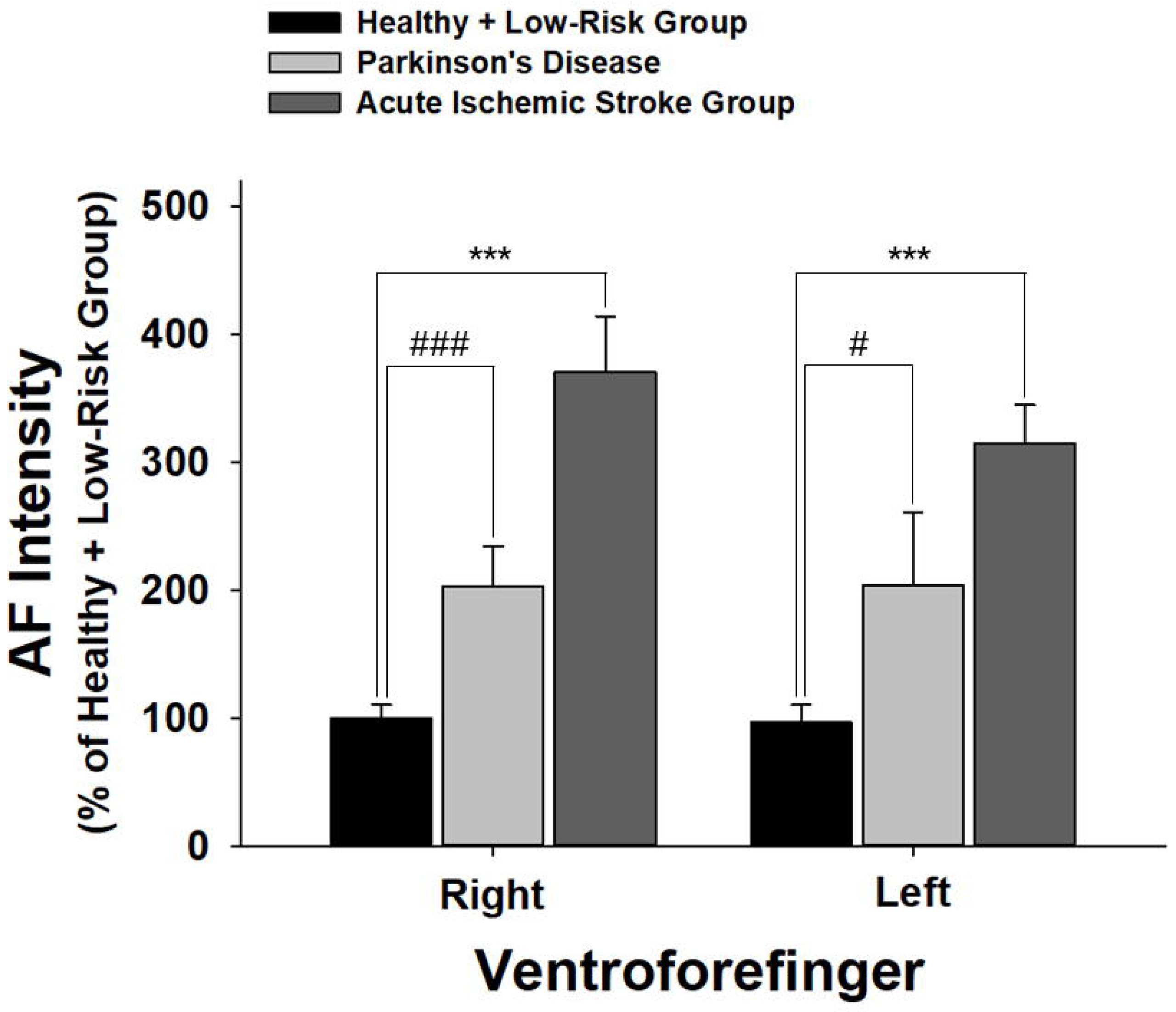
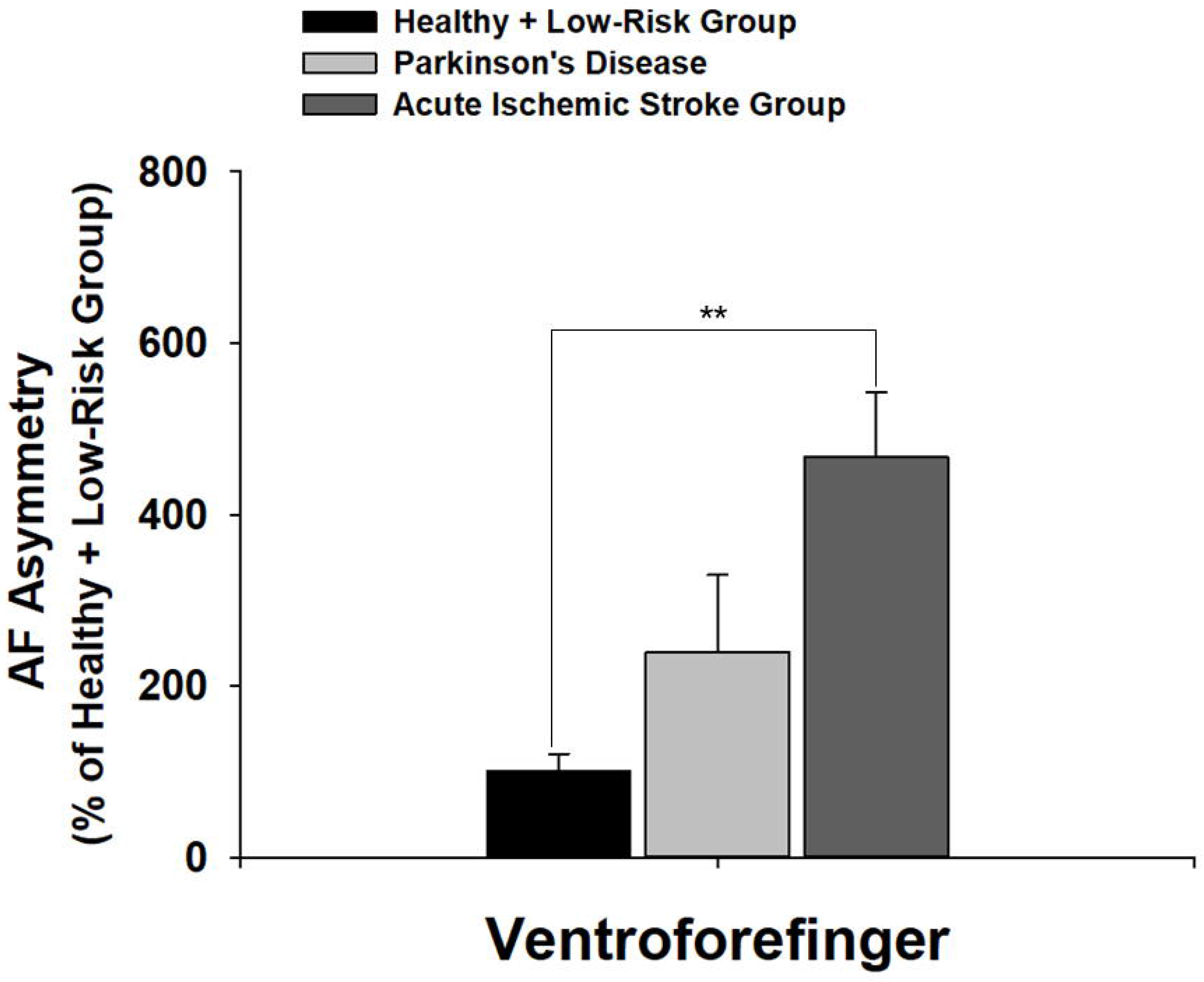
(A) On right and left Ventroforefinger, both PD patients and AIS patients have significantly higher green AF intensity than Healthy and Low-Risk persons. (B) The AIS patients have significantly higher AF asymmetry than Healthy and Low-Risk persons at the Ventroforefingers. The number of subjects in the Healthy and Low-Risk group, the PD group, and the AIS group is 25, 13, and 47-49, respectively. **, *p* < 0.01; ***, *p* < 0.#, *p* < 0.05 (Student *t*-test); ###, *p* < 0.001 (Student *t*-test).

We further found that on both right and left side of Ventriantebrachium, PD patients have significantly higher green AF intensity than Healthy and Low-Risk persons (Fig. 3A), while AIS patients have significantly higher green AF intensity than Healthy and Low-Risk persons only at right Ventriantebrachium (Fig. 3A). Both PD patients and AIS patients also have significantly higher AF asymmetry than Healthy and Low-Risk persons (Fig. 3B). We also found that on left Index Fingernail, PD patients have significantly higher green AF intensity than Healthy and Low-Risk persons (Fig. 4A), while AIS patients have significantly higher green AF intensity than Healthy and Low-Risk persons on both left and right Index Fingernails (Fig. 4A). Both PD patients and AIS patients also have significantly higher AF asymmetry than Healthy and Low-Risk persons (Fig. 4B).

**Fig. 3.**
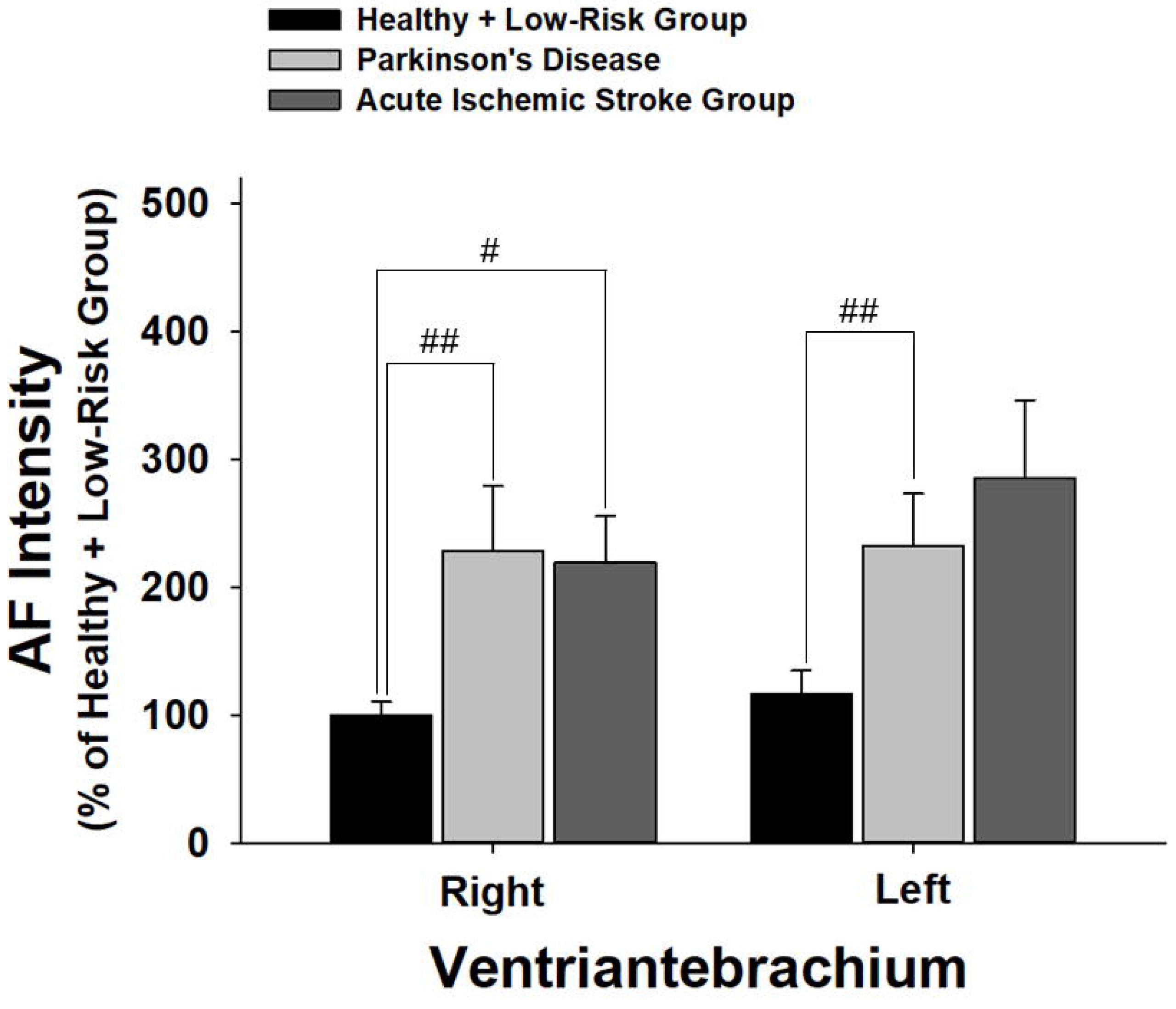
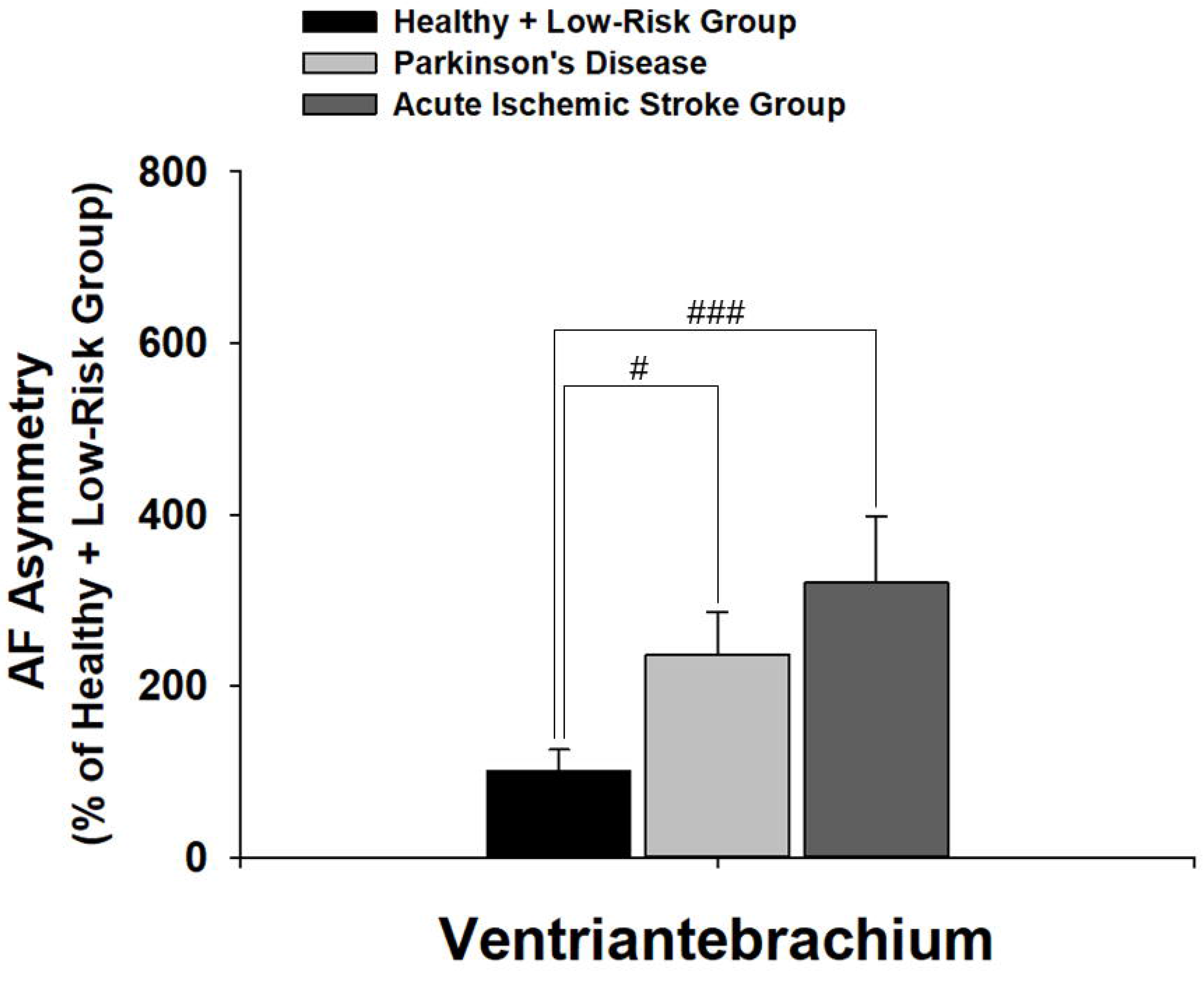
(A) On both right and left Ventriantebrachium, PD patients have significantly
higher green AF intensity than Healthy and Low-Risk persons, while AIS patients have significantly higher green AF intensity than Healthy and Low-Risk persons only at right Ventriantebrachium. (B) Both PD patients and AIS patients have significantly higher AF asymmetry than Healthy and Low-Risk persons at the Ventriantebrachium. The number of subjects in the Healthy and Low-Risk group, the PD group, and the AIS group is 25, 13, and 47-49, respectively. #, *p* < 0.05 (Student *t*-test); ##, *p* < 0.01 (Student *t*-test); ###, *p* < 0.001 (Student *t*-test).

**Fig. 4.**
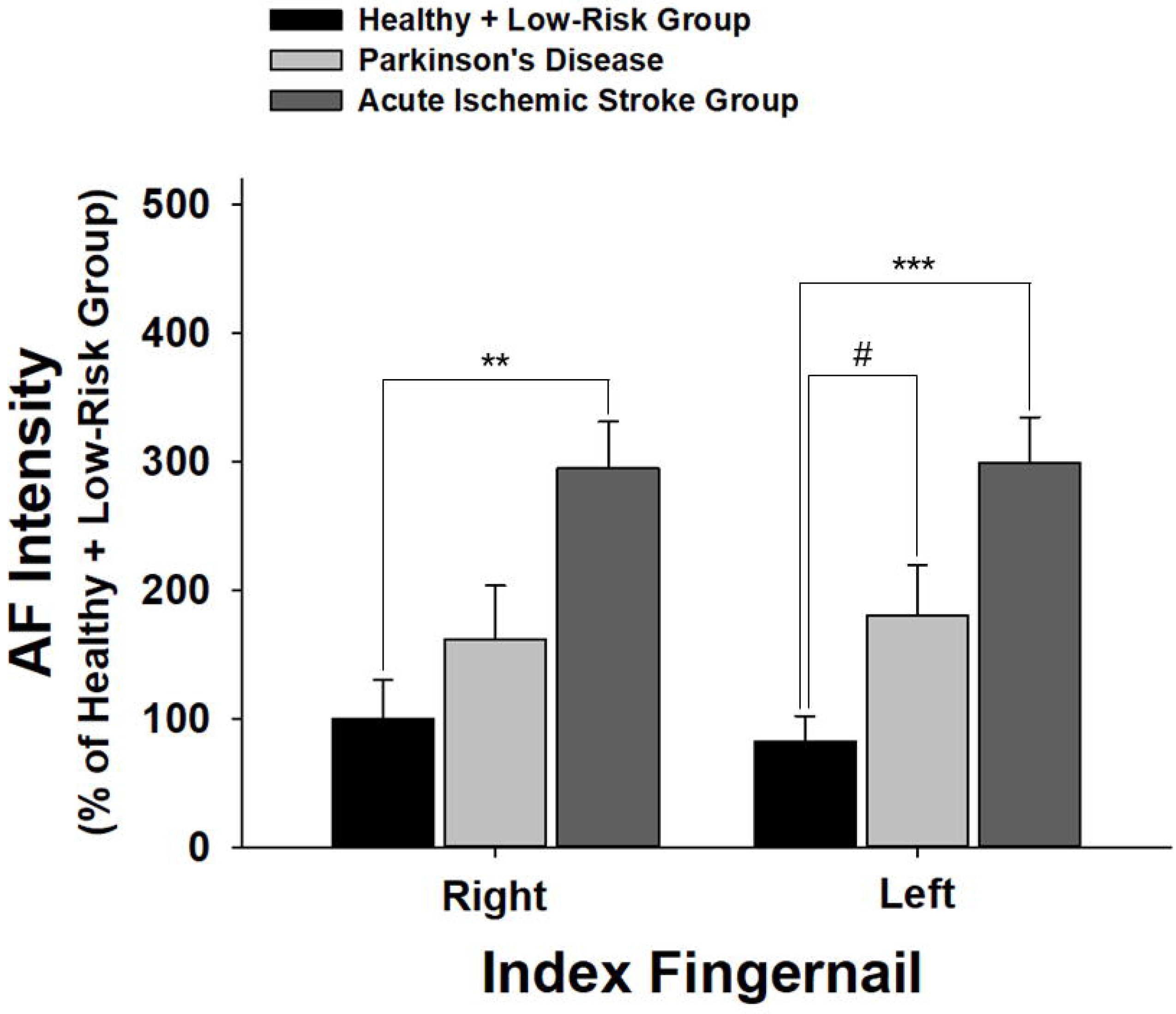
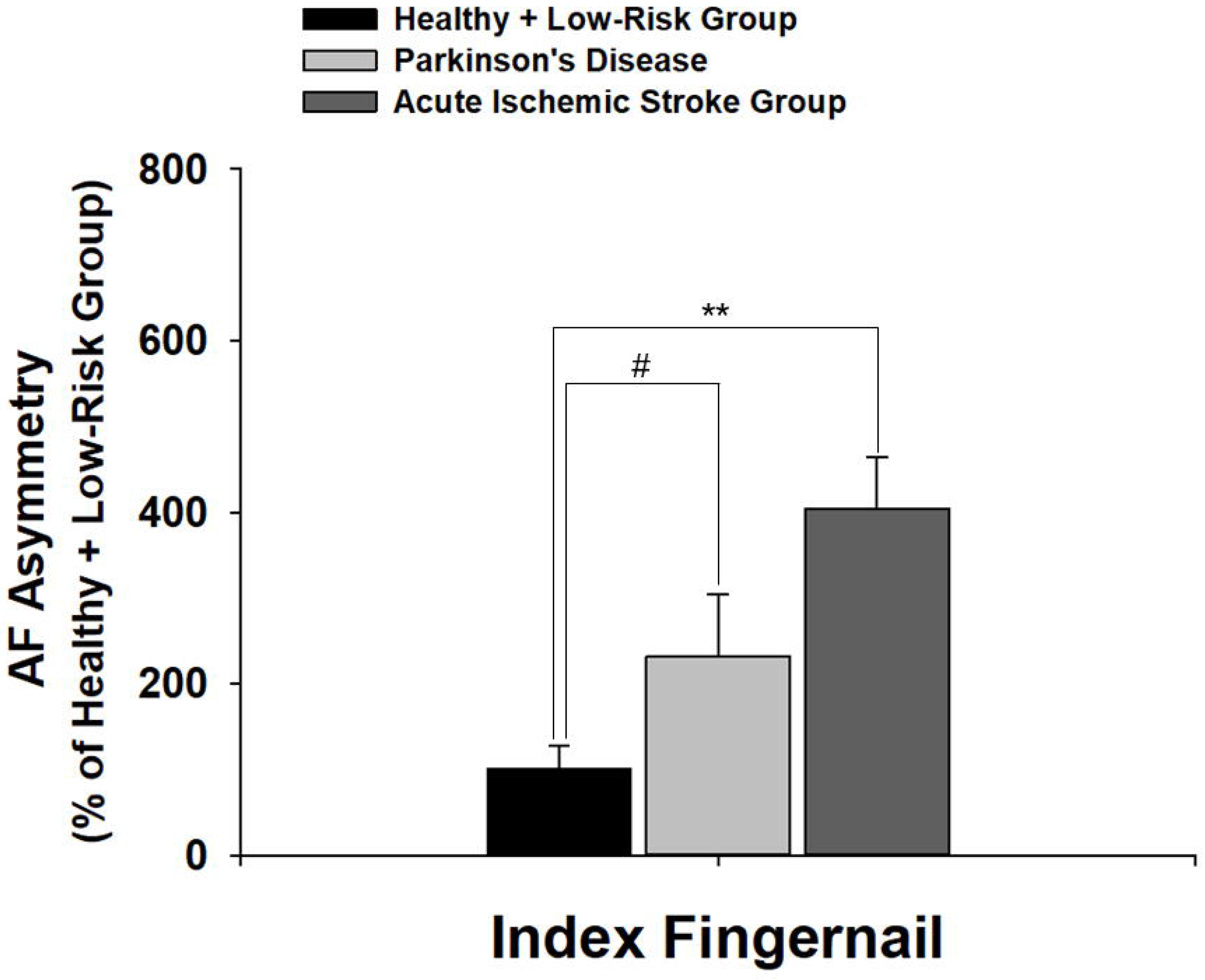
(A) On left Index Fingernail, PD patients have significantly higher green AF intensity than Healthy and Low-Risk persons, while AIS patients have significantly higher green AF intensity than Healthy and Low-Risk persons on both left and right Index Fingernails. (B) Both PD patients and AIS patients have significantly higher AF asymmetry than Healthy and Low-Risk persons at Index Fingernails. The number of subjects in the Healthy and Low-Risk group, the PD group, and the AIS group is 25, 13, and 47-49, respectively. **, *p* < 0.01; ***, *p* < 0.001; #, *p* < 0.05 (Student *t*-test).

On the right dorsal side of index fingers, PD patients have significantly higher green AF intensity than Healthy and Low-Risk persons (Fig. 5A), while AIS patients have significantly higher green AF intensity than Healthy and Low-Risk persons on both left and right dorsal side of index fingers (Fig. 5A). Only AIS patients, but not PD patients, have significantly higher AF asymmetry than Healthy and Low-Risk persons (Fig. 5B). Our study also showed that on both right and left Centremetacarpus, AIS patients have significantly higher green AF intensity than PD patients as well as Healthy and Low-Risk persons, while there is no difference in the AF intensity between the PD group and the Healthy and Low-Risk group (Fig. 6A). AIS patients also have significantly higher AF asymmetry than PD patients as well as Healthy and Low-Risk persons, while there is no difference in the AF asymmetry between the PD group and the Healthy and Low-Risk group (Fig. 6B).

**Fig. 5.**
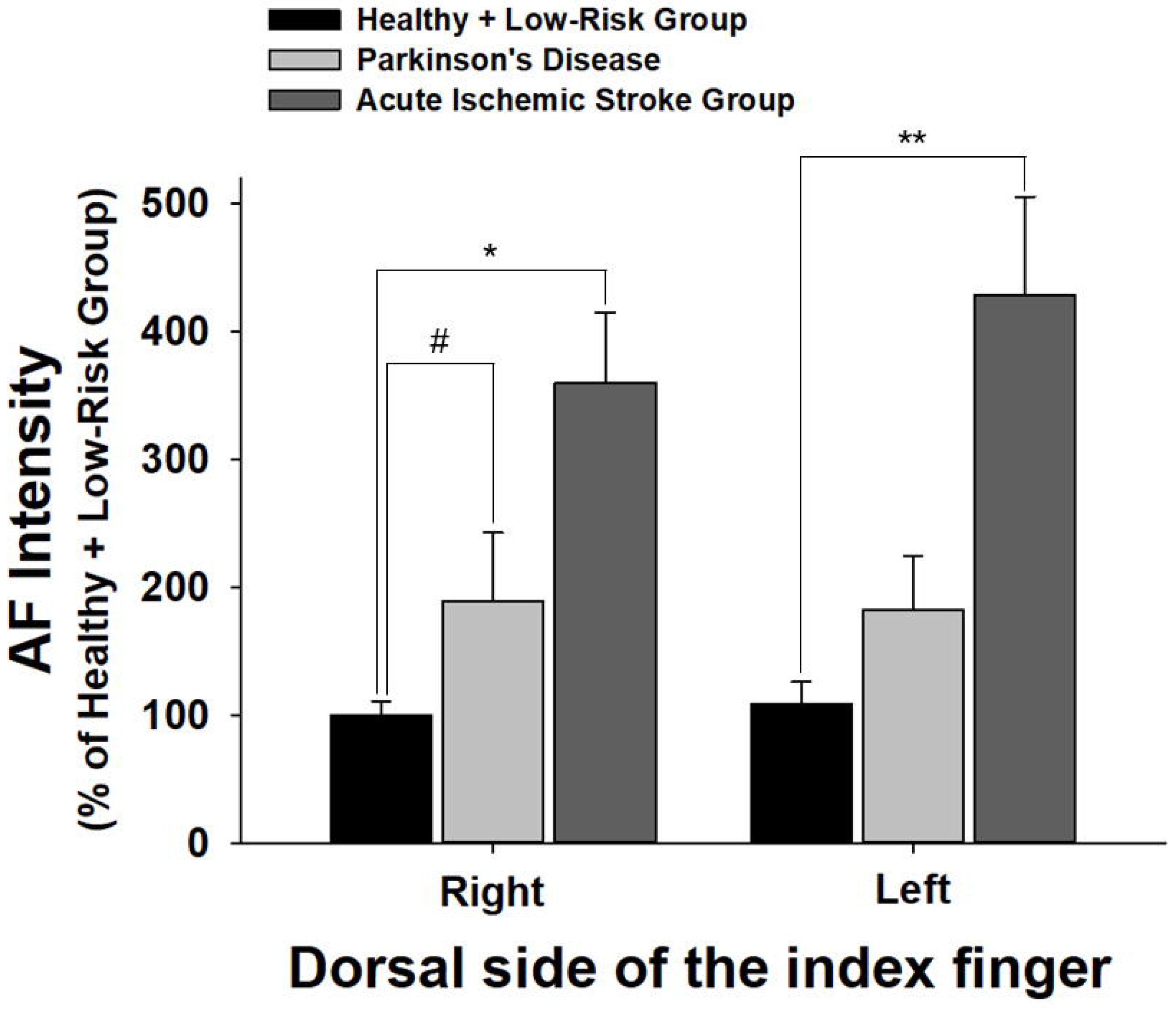
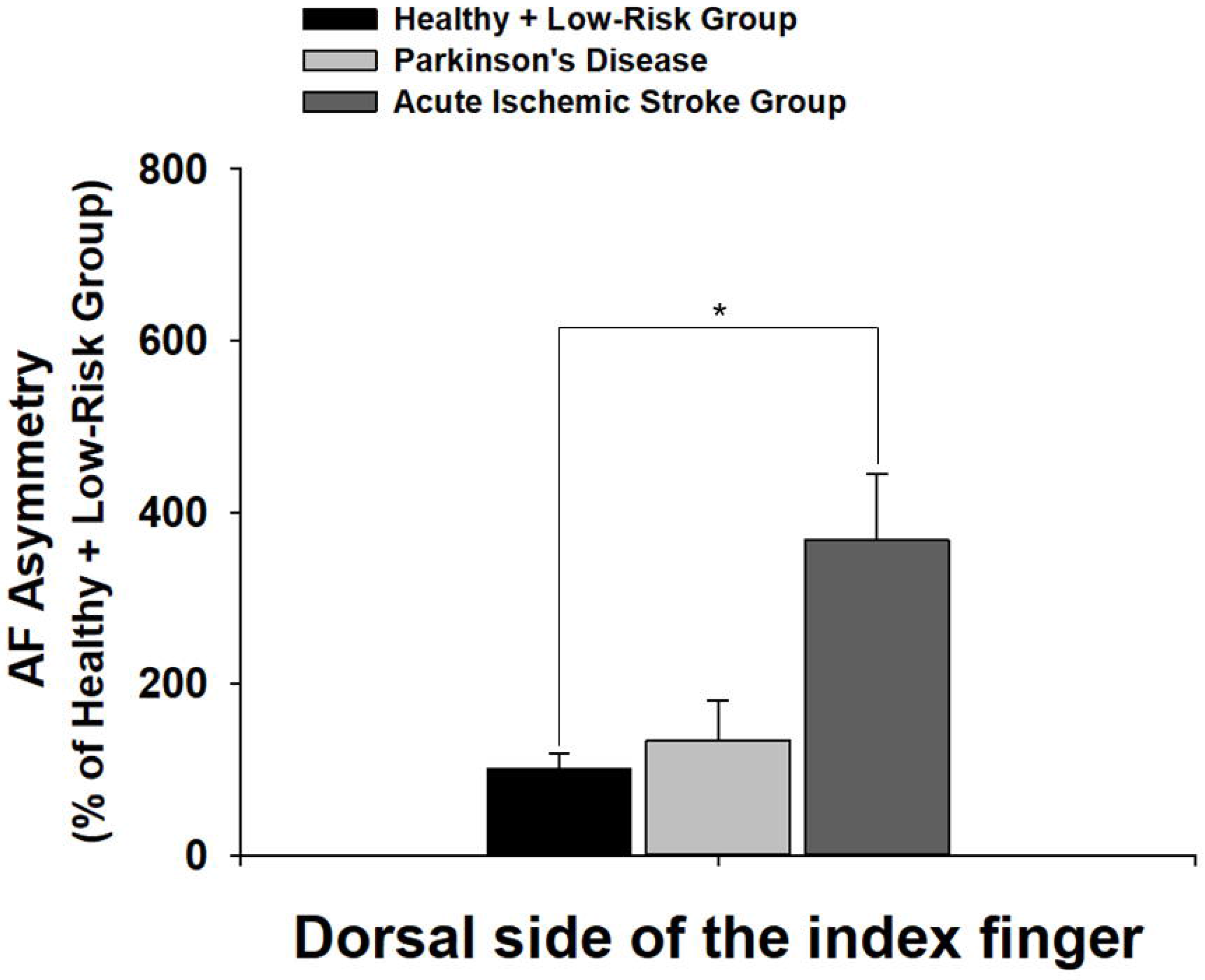
(A) On the right dorsal side of index finger, PD patients have significantly higher green AF intensity than Healthy and Low-Risk persons, while AIS patients have significantly higher green AF intensity than Healthy and Low-Risk persons on both left and right dorsal side of index finger. (B) Only AIS patients, but not PD patients, have significantly higher AF asymmetry than Healthy and Low-Risk persons at dorsal side of index finger. The number of subjects in the Healthy and Low-Risk group, the PD group, and the AIS group is 25, 13, and 47-49, respectively. *, *p* < 0.05; **, *p* < 0.01; #, *p* < 0.05 (Student *t*-test).

**Fig. 6.**
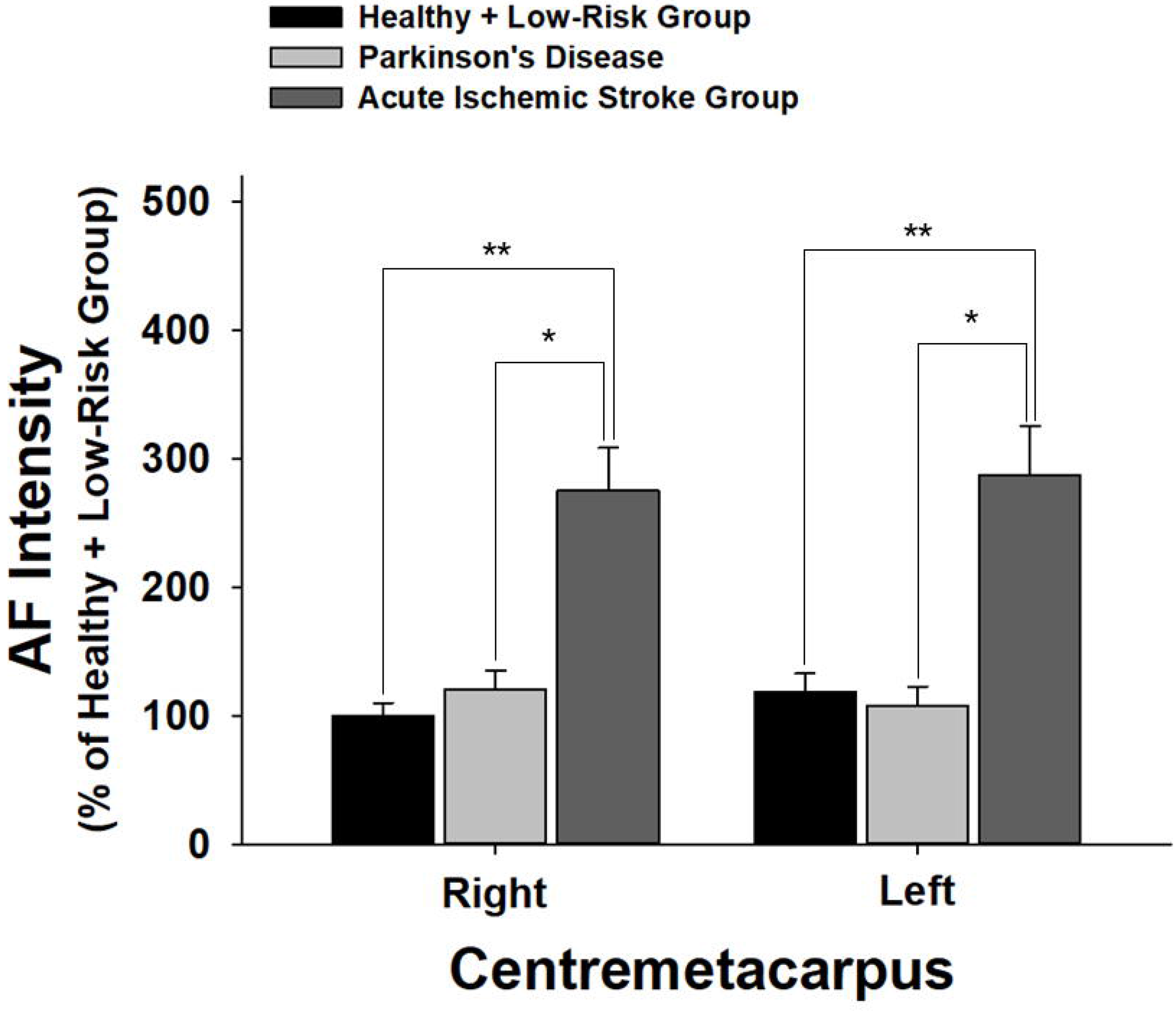
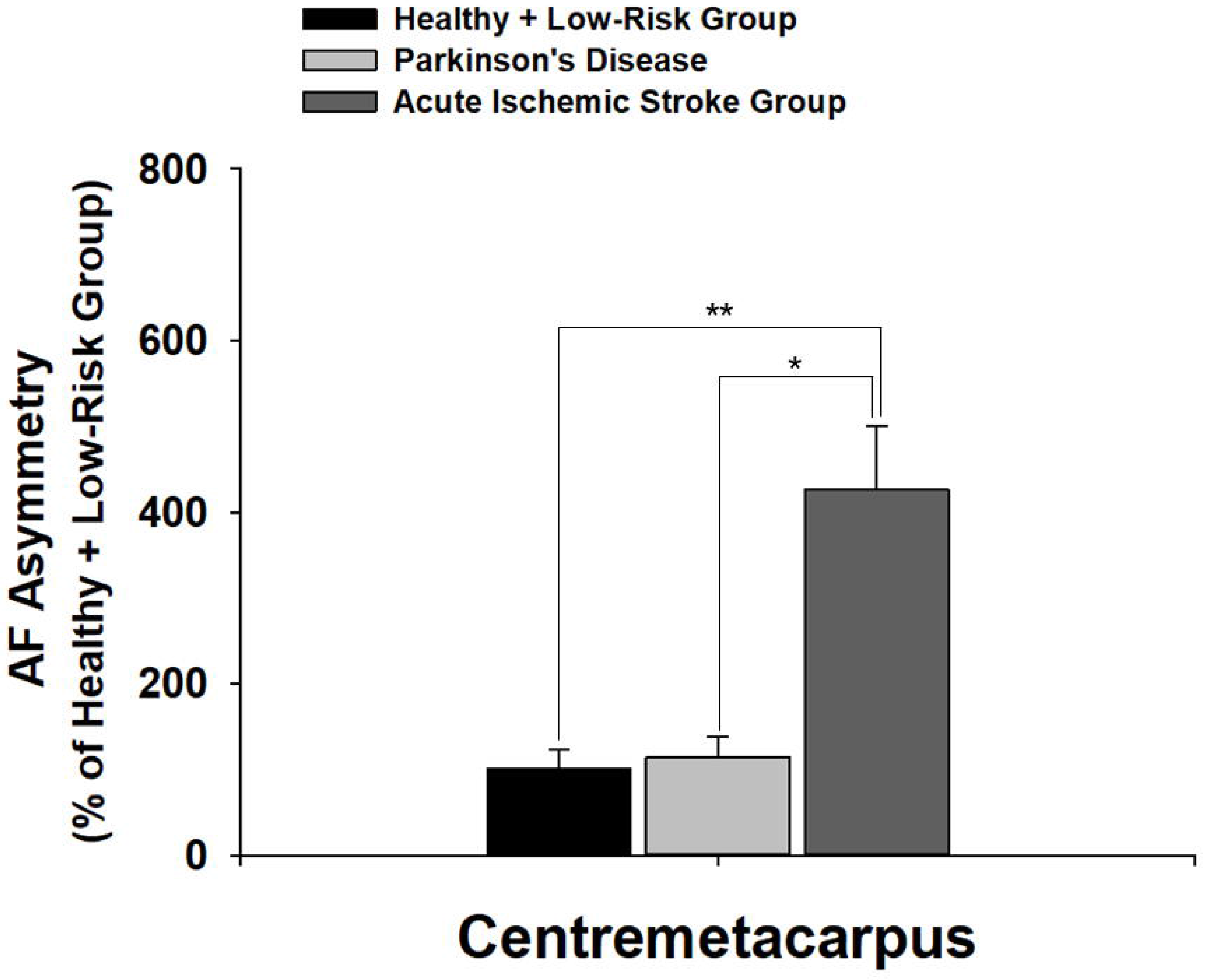
On both right and left Centremetacarpus, AIS patients have significantly higher green AF intensity than PD patients as well as Healthy and Low-Risk persons, while there is no difference in the AF intensity between the PD group and the Healthy and Low-Risk group. (B) AIS patients have significantly higher AF asymmetry than PD patients as well as Healthy and Low-Risk persons at Centremetacarpus, while there is no difference in the AF asymmetry between the PD group and the Healthy and Low-Risk group at Centremetacarpus. The number of subjects in the Healthy and Low-Risk group, the PD group, and the AIS group is 25, 13, and 47-49, respectively. *, *p* < 0.05; **, *p* < 0.01.

## Discussion

Our current study has indicated that PD patients have a distinct pattern of AF changes, compared with those of healthy and Low-Risk persons as well as AIS patients: First, the AF intensity of PD patients in the fingernails and a part of the regions of examined skin is significantly higher than that of the healthy and Low-Risk group, while in most regions examined the AF intensity of AIS patients is significantly higher than that of the healthy and Low-Risk group; second, there is AF asymmetry at index fingernails and two regions of the skin of PD patients, while there is AF asymmetry at all examined regions of AIS patients; and third, both the AF intensity and AF asymmetry at Centremetacarpus of PD patients is significantly lower than those of AIS patients. Moreover, it appears that the average levels of the increases in the AF intensity of PD patients are markedly lower than those of AIS patients. Collectively, our study has indicated that PD patients have a distinct pattern of AF changes compared with those of healthy and Low-Risk persons as well as AIS patients. This PD patients’ distinct pattern of AF changes may become a novel diagnostic biomarker for PD.

Our current study has provided the first evidence suggesting that PD patients may be diagnosed at homes by determining the AF of fingernails and certain regions of their skin. Based on the evidence shown in our study, we proposed a new criteria for PD diagnosis - ‘selective AF increases in certain regions of skin’, which may be used jointly with the clinical symptom-based diagnostic criteria for future PD diagnosis. The sensitivity and specificity of this diagnostic approach may be significantly enhanced in the future, with increases in both the regions examined and applications of deep machine learning and big data technology.

Our previous study has shown that the oxidative stress induced by UVC mediates the increase in the epidermal green AF of mouse ears by inducing keratin 1 proteolysis (10). Since there are significant increases in oxidative stress in the plasma of PD patients (2,4,6), we propose that the increased oxidative stress may induce increases in the epidermal AF of the PD patients by inducing keratin 1 proteolysis.

As discussed above, there are at least three major problems and challenges in the current protocols for PD diagnosis. Our novel AF-based approach may significantly enhance our capacity to overcome these problems and challenges: First, nonspecific clinical symptoms of patients constitute fundamental basis of current PD diagnosis (1), which belongs to semi-quantifiable or non-quantifiable information. The accuracy and efficacy of PD diagnosis has been limited by the lack of biomarker tests (1). Our AF-based method may become a biomarker test for PD, which belongs to quantifiable information that may significantly increase the accuracy and efficacy of PD diagnosis. Second, dependence on the clinical judgment of experienced neurologists has also been a major problem in PD diagnosis, due to the shortage of experienced neurologists in many areas. This problem may be significantly alleviated by our AF-based diagnostic approach, which may transform the current experience-based diagnostic approach into a medical device-based approach. Third, the accuracy of diagnosis for early-stage PD has been highly unsatisfactory (1). Our AF-based diagnostic approach provides quantifiable information. This information, together with the information of clinical symptoms of patients, could be analyzed by AI technology, which may lead to significantly increased accuracy of diagnosis for early-stage PD.

## Acknowledgment

The authors would like to acknowledge the financial support by Major Research Grants from the Scientific Committee of Shanghai Municipality #16JC1400500 and #16JC1400502 (to W.Y.) and Chinese National Natural Science Foundation Grant #81271305 (to W. Y.).

